# *GTP cyclohydrolase II* (*gch2*) and axanthism in ball pythons: a new vertebrate model for pterin-based pigmentation

**DOI:** 10.1101/2024.05.22.595308

**Authors:** Alan Garcia-Elfring, Heather L. Roffey, Jaren M. Abergas, Andrew P. Hendry, Rowan D. H. Barrett

## Abstract

Pterin pigments are responsible for many of the bright colours observed across the animal kingdom. However, unlike melanin, the genetics of pterin-based pigmentation has received relatively little attention in animal colouration studies. Here, we investigate a lineage of axanthic ball pythons (*Python regius*) found in captivity as a model system to study pterin pigmentation in vertebrates. By crowdsourcing shed skin samples from commercial breeders and applying a case-control study design, we utilized whole-genome pool sequencing (pool-seq) and variant annotation. We identified a premature stop codon in the gene *GTP cyclohydrolase II* (*gch2*), which is associated with the axanthic phenotype. GCH2 catalyzes the first rate-limiting step in riboflavin biosynthesis. This study provides the first identification of an axanthism-associated gene in vertebrates and highlights the utility of ball pythons as a model to study pterin-based pigmentation.

## INTRODUCTION

Animal pigmentation has long served as a valuable model for understanding genetics, development, and evolution (Cott 1940; Kronforst et al. 2012; Caro 2017; Endler et al. 2017; San-Jose et al. 2017). Vertebrate pigmentation arises from the differential absorption of light by pigments within specialized cells called melanophores (melanocytes in endothermic vertebrates) and xanthophores. All vertebrates produce a brown/black form of melanin derived from the amino acid tyrosine (Wakamatsu and Ito 2021). In addition, poikilothermic vertebrates (fish, amphibians, and reptiles) display bright colouration through yellow/red pigments found in cells called xanthophores. Xanthophores derive their vibrant yellow or red colouration from pterin or carotenoid pigments. While vertebrates acquire carotenoids exclusively from their diet (Maoka 2020), pterin pigments are synthesized de novo from guanosine triphosphate (GTP). Coloration can also be structural, with poikilothermic vertebrates using guanine nanocrystals in iridophore cells to generate structural colouration via the constructive interference of light. Although melanin-based pigmentation has been extensively studied, other pigment classes, such as pterins, have received less attention.

The mouse has been pivotal in studies of melanocyte biology, helping uncover the genes responsible for melanocyte development and patterning: *Kit*, *Pax3*, *Mitf*, *Edn3*, *Ednrb*, and *Sox10* (Steingrímsson et al. 2005); melanin synthesis: *Tyr*, *Tyrp1*, *Dct* (McNamara et al. 2021; Wakamatsu and Ito 2021); and its regulation, such as *Mc1r* (Garcia-Borron et al. 2005). Meanwhile, the zebrafish model has laid the foundation for understanding poikilothermic vertebrate pattern formation (Singh and Nüsslein-Volhard 2015). Studies on zebrafish have shown that pattern formation arises from local, short- and long-range cell-cell interactions between chromatophore types (Frohnhöfer et al. 2013). For example, zebrafish mutants lacking any one of the chromatophores (iridophores, melanophores, and xanthophores) fail to develop stripes and often develop spots instead. In double mutants lacking two of the three chromatophore types, the remaining chromatophores uniformly cover the body (Frohnhöfer et al. 2013; Owen et al. 2020). Gap junction proteins mediate these cell-cell interactions, and mutations in these genes can result in labyrinthine patterning (Irion et al. 2014). However, it remains unclear how widely zebrafish cell-cell interactions apply across vertebrate taxa.

First discovered in butterfly wings (Hopkins 1895), pterin-based pigmentation is ubiquitous in nature but has received less attention compared to melanin-based pigmentation (Andrade and Carneiro 2021). The pterin pigment synthesis pathway has been largely elucidated through studies in *Drosophila* (Glassman and Mitchel 1959; Courtright 1967; Ziegler and Harmsen 1970; Goto and Sugiura 1971; Evans and Howells 1978; Spradling and Rubin 1983; Reaume et al. 1991; Kim et al. 2013). Notably, *Drosophila*’s first described mutant, “white,” lacked eye pterin (drosopterin) and ommochrome pigments (Morgan 1910). While melanin is often linked to adaptive phenotypes, such as cryptic colouration (Barrett et al. 2019; Harris et al. 2020; Baltazar-Soares et al. 2024), pterin pigments are key in signaling both within and between species (Kikuchi and Pfennig 2012; Kikuchi et al. 2014; Andrade and Carneiro 2021). In vertebrates, zebrafish have been the primary model for studying xanthophore biology, including xanthophore specification from neural crest cells (Kelsh et al. 1996; Odenthal et al. 1996; Parichy et al. 2000; Minchin and Hughes 2008) and pigment synthesis (Ziegler 2003; Lister 2019).

Studying mutants lacking a pigment can offer insights into bright colouration genetics, a strategy pioneered by Thomas Hunt Morgan with fruit flies. Axanthic animals, which lack yellow and red pigments but retain melanin, are rare among vertebrates. Only a few axanthic amphibians and reptiles have been discovered in nature (Bechtel 1991; Jablonski et al. 2014; Cattaneo 2015; Kolenda et al. 2017; Cavalcante and Bruni 2018; Borteiro et al. 2021; Schluckebier et al. 2022). Axanthic vertebrate models are also scarce in pigmentation research, with the axanthic axolotl, first described by Lyerla and Dalton (1971), being a notable exception (Dunson 1974; Frost et al. 1986; Bukowski et al. 1990; Masselink and Tanaka 2021). Axanthic axolotls lack both xanthophores and iridophores, meaning axanthism in axolotls affects chromatophore development rather than pigment synthesis. However, low survivability in axanthic axolotls presents challenges for research, including breeding programs, population genetics studies, and comparative analyses.

Here, we use a line of axanthic ball pythons to study the genetics of yellow pigmentation in reptiles. We hypothesize that, assuming a similar mechanism of vertebrate pattern formation to zebrafish (Frohnhöfer 2013; Singh and Nüsslein-Volhard 2015), axanthism in ball pythons results from defects in pterin pigment synthesis (Ziegler 2003), rather than carotenoid metabolism or xanthophore development. We expect that xanthophore development remains unaffected, as axanthic ball pythons exhibit wild-type melanin patterning, which suggests intact cell-cell interactions. Furthermore, we anticipate that yellow pigmentation (or its absence) is linked to pterin-based pigmentation, rather than carotenoids, since ball pythons are born with yellow colouration before acquiring carotenoids from their diet. This suggests that pterins, not carotenoids, underlie the colouration difference.

## METHODS

### Sample collection, study design, DNA extraction, and sequencing

We crowd-sourced ball python shed skin samples from breeders, including five samples with the recessive VPI Axanthic colour morph (Table S1). VPI refers to Vida Preciosa International, the reptile propagation company that first received the wild-caught axanthic snake from Western Africa and established the phenotype in captivity. All VPI Axanthic (hereafter referred to as ‘axanthic’) ball pythons descend from this individual.

Additionally, we collected 38 samples of the recessive ‘Clown’ phenotype (Table S2), which we used as a control for identifying candidate mutations linked to the axanthic trait. We also gathered samples from other colour morphs: ‘Spider’ (n=26, Table S3), ‘Enchi’ (n=7, Table S4), ‘Ivory’ (n=15, Table S5), and ‘Lavender’ (n=14, Table S6). These were used as further control sets because, according to breeder pedigrees, these morphs do not carry any axanthism variation.

Similarly, the axanthic samples were confirmed to lack any variation associated with the Clown, Spider, Enchi, Ivory, or Lavender morphs (i.e., inferred to be homozygous for the reference alleles at those loci). We identified candidate mutations by comparing genetic variation in the axanthic pool with multiple reference sample sets known, through pedigree analysis, to not carry the axanthic variant.

The wild-type ball python colour morph displays a black background with tan-coloured lateral saddles and traces of yellow pigmentation (Figure 1A). Comparing the wild type to the albino colour morph highlights normal xanthophore pigmentation and patterning: in non-axanthic individuals, yellow pigment typically overlaps with the tan-coloured saddles and is absent from the dark melanic regions (Figure 1B). In contrast, axanthic ball pythons lack yellow pigmentation but retain the wild-type melanin pattern (Figure 1C).

**Figure 1.**
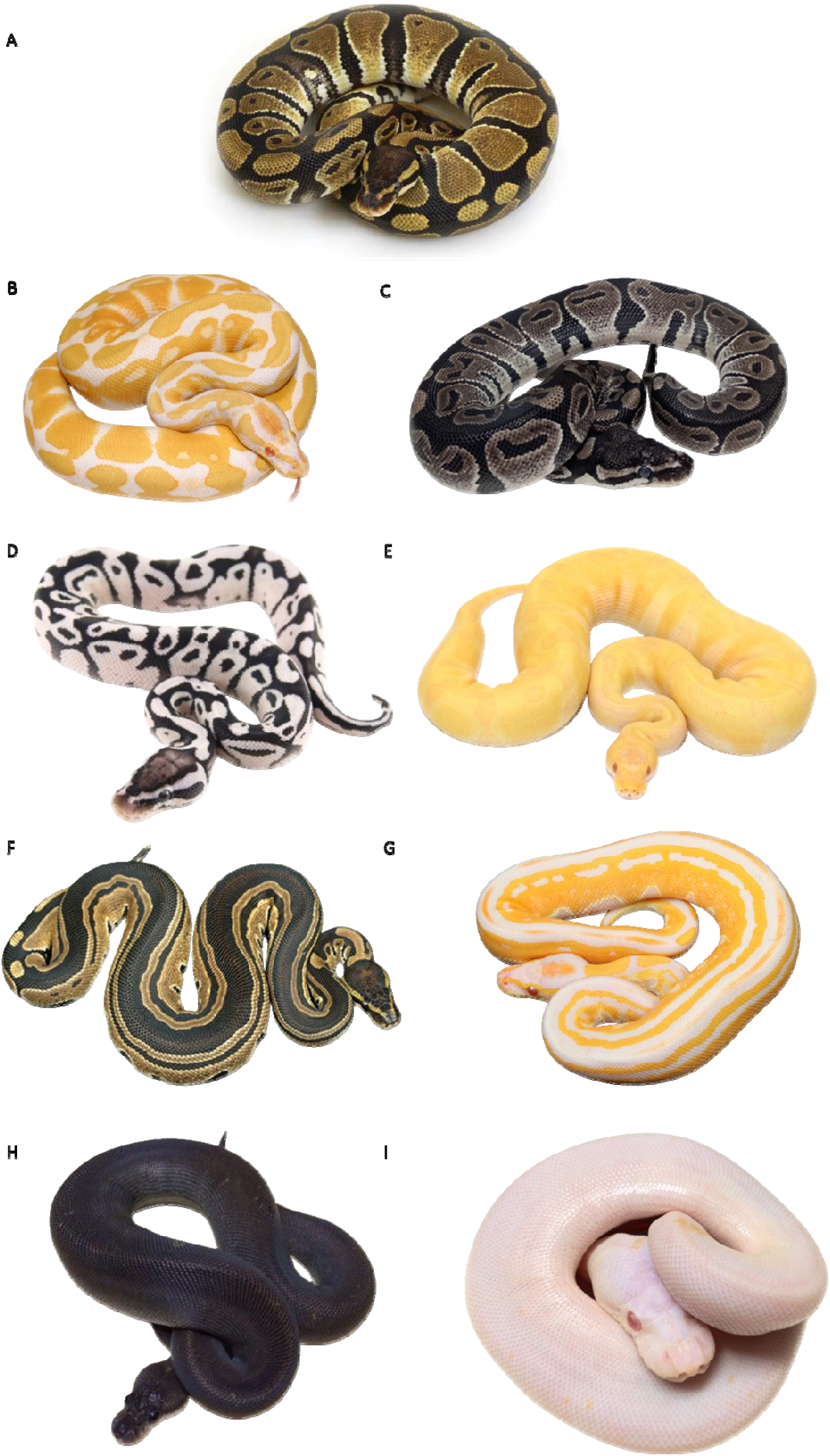
Mendelian phenotypes (colour morphs) in ball pythons. Compared to the (A) wild type, (B) the albino colour morph lacks melanin pigment which allows the display of xanthophore (pterin) pigmentation and patterning, occupying the tan-coloured parts of the wild type. (C) The VPI Axanthic is characterized by a lack of yellow pigmentation on a wildtype pattern. (D) Combining colour morphs with axanthism (VPI Axanthic + Pastel + Fire is shown) can create different forms of the colour anomaly. (E) Some colour morphs like Enchi (Albino + Enchi is shown) are associated with changes in xanthophore migration (e.g., xanthism). (F) Tri-Stripe colour morph exhibits altered melanin patterning. (G) The Albino form of Tri-Stripe shows the effects of this colour morph on xanthophore pigmentation. (H) Melanic colour morphs like Cinnamon do not have xanthophore pigmentation, as shown by an (I) all-white snake in the Albino form. Photo credit: worldofballpythons.com

We extracted and sequenced DNA from the shed skins using a standard three-day phenol-chloroform procedure. DNA quantification and quality checks were performed with an Infinite® 200 Nanoquant (Tecan Group Ltd., Männedorf, Switzerland). After normalizing the DNA from each sample, we combined them into pools in equimolar amounts. These pooled samples (case and controls) were then sent to the McGill Genome Centre for whole-genome sequencing. Library preparation was performed using a PCR-based method, and sequencing was carried out with 150 bp paired-end reads on one lane of the Illumina NovaSeq 6000 platform.

### Bioinformatics

We used the trim-fastq.pl program from Popoolation (Kofler et al. 2011a) to filter raw reads based on quality (--quality-threshold 20) and length (--min-length 50). This step ensured that each read had a minimum quality score of 20 and a minimum length of 50 bp. We then mapped the filtered reads to a reference genome using the bwa-mem program with default parameters (Li and Durbin, 2009). Since a ball python reference genome is not available, we aligned the reads to both the draft Burmese python reference genome *Pmo2.0* (Castoe et al. 2013) and the more recent chromosome-length Burmese python assembly (*Python molurus bivittatus*-5.0.2_HiC), available at DNAzoo.org. This chromosome-length assembly, based on Hi-C data (Dudchenko et al. 2017; Dudchenko et al. 2018), is aligned to the annotated draft genome (Castoe et al. 2013) and has been used in previous studies to identify colour-related loci (Garcia-Elfring et al. 2023).

Next, we converted SAM files to BAM format using SAMtools (Li et al. 2009) and removed reads with a mapping quality below 20 (-q 20). We then generated mpileup and sync formats using SAMtools and Popoolation2 (Kofler et al. 2011b), respectively, before proceeding to variant calling. Variants were called and filtered with bcftools (vcfutils.pl varFilter) using a minimum depth of five (-d 5) and a mapping quality of 20 (-Q 20).

We calculated SNP-specific F_ST_ estimates using fst-sliding.pl from Popoolation2 (Kofler et al. 2011b). F_ST_ values were generated for SNP variants in VPI Axanthic (n=5) compared to the control pools. Since the control pools (e.g., Clown, Spider, etc.) lacked axanthic variation, we expected the derived allele to be fixed in the axanthic samples and the reference allele to be fixed in the control pool. A threshold F_ST_ value of 1.0 was used to identify candidate mutations. To confirm the genomic region and candidate mutation of interest, we generated F_ST_ values between the axanthic samples and additional control pools (Spider, n=26; Enchi, n=7; Ivory, n=15; Lavender, n=14).

### Variant annotation and identification of polymorphisms and fixed differences

We used the software SnpEff (Cingolani et al. 2012) to annotate variants mapped to both the draft and chromosome-length assemblies. SnpEff utilizes a genome assembly and its corresponding General Feature Format (GFF3) file to assess the functional impact of variants on genomic features. For genomic regions showing elevated FST values (“F_ST_ peaks”), we focused on individual variants with an F_ST_ of 1.0, that mapped to protein-coding regions and were predicted to affect gene products. These included nonsense mutations and small indels classified as “high impact,” and missense mutations classified as “moderate impact,” which were considered candidate mutations.

To visualize the position and effect of a candidate mutation on the protein, we used the program Protter (Omasits et al. 2016). We confirmed the candidate variant by PCR amplification and Sanger sequencing of the locus using samples that were not included in the pooled libraries. We validated the candidate loci by Sanger sequencing samples with genotypes inferred from breeder pedigrees. These included inferred heterozygotes (n=10), homozygotes for the derived axanthism-causing allele (i.e., axanthic individuals, n=6), and inferred homozygotes for the wild-type allele (n=10).

Primers used to amplify the region surrounding the candidate locus were designed using Primer3 (Kõressaar and Remm, 2007; Kõressaar et al. 2018). The primers used for PCR amplification were as follows:

- Forward: 5’- GAAACTTTTGACCAGACTCAGTC
- Reverse: 5’- CTTCAATCACCACAGCCACTCC

### Synteny and phylogenetic analysis of gch1 and gch2

We conducted a synteny analysis and constructed a phylogenetic tree for the *gch1* and *gch2* genes across multiple species: Burmese python (*Python bivittatus*), brown anole (*Anolis sagrei*), coelacanth (*Latimeria chalumnae*), and the elephant shark (*Callorhinchus milii*). To ensure focus on conserved regions, we trimmed the gene sequences (555 bp) before alignment. Due to the evolutionary divergence of these species, we used MAFFT to build the tree(Katoh et al. 2002). Phylogenetic trees were then built using FastTree (Price et al. 2010), which estimates maximum-likelihood phylogenies, allowing us to infer the evolutionary relationships of these genes across species.

## RESULTS

Our whole-genome sequencing and variant calling identified 26,449,779 SNPs across all samples received from ball python breeders. Analysis of allele frequency differences between the axanthic pool and the control pool (Clown colour morph) demonstrates overall high levels of background F_ST_, likely resulting from the low sample size for the axanthic phenotype (n=5). We find two areas of high differentiation or F_ST_ peaks. One region (∼24.8 Mb) on chromosome two (Figure 2A) and a second peak on chromosome three, corresponding to the region containing the Clown mutation (Garcia-Elfring et al. *in prep*). Comparing the axanthic data with additional control pools (Spider, n=26; Enchi, n=7; Ivory, n=15; Lavender, n=14) confirms that the region of high differentiation on chromosome two is associated with axanthism (Figure S1). Within the peak on chromosome two, one variant had both high levels of differentiation (F_ST_ = 1) and is predicted to have a high impact on the gene product. This variant is a nonsense mutation, and it maps to exon 5 of LOC103050242 (*GTP cyclohydrolase 1-like*) in the draft assembly. The candidate SNP consists of a C to T nucleotide change (c.520C>T), changing an arginine residue (CGA) to a TGA stop codon (p.Arg174*). This genetic variant is expected to result in a truncated protein (Figure 2B). We validated the axanthic variant found in poolseq data by PCR amplification and Sanger sequencing of the locus of interest in additional individuals (inferred heterozygotes: n=10; homozygotes for the derived axanthism-causing allele: n=6; inferred homozygotes for the wild type allele: n=10). All individuals tested had the genotype that was inferred from pedigrees by breeders.

**Figure 2.**
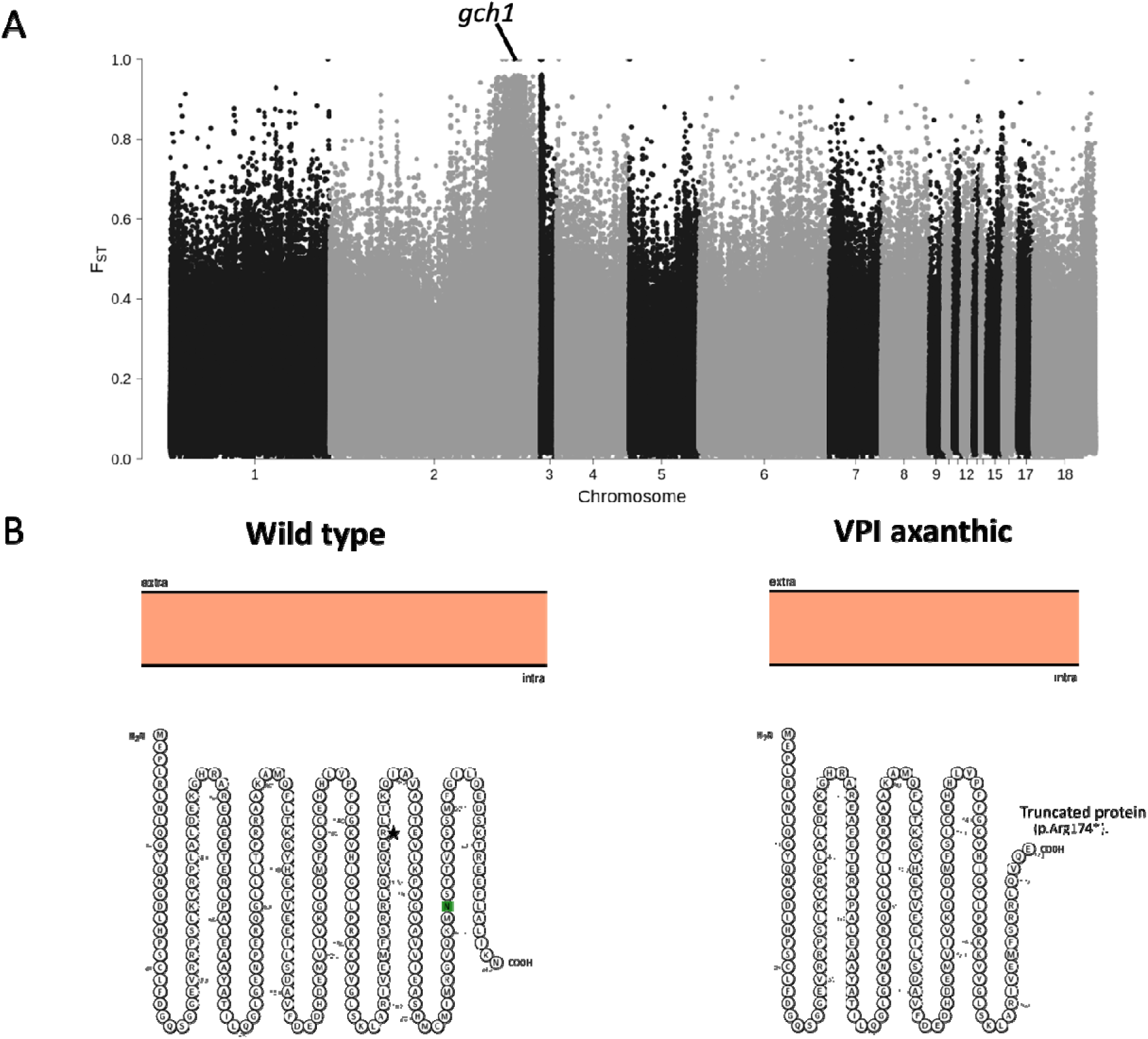
(A) Manhattan plot of F_ST_ values for SNPs mapped to the chromosome-length assembly. Genomic differentiation between VPI axanthic and a reference pool (Clown colour morph) shows a peak on chromosome two housing the axanthism candidate variant on LOC103050242 (*gch2*). (B) Burmese python amino acid sequence of LOC103050242, with the VPI Axanthic locus indicated by an asterisk (left panel). A nonsense mutation the on the 174^th^ residue (p.Arg174*) likely leads to a defective truncated protein (right panel) and axanthism.

Based on the annotation of *gch1* in the Burmese python reference genome and the absence of an annotated *gch2*, we hypothesized that the gene LOC103050242 (*GTP cyclohydrolase 1-like*) corresponds to the paralog *gch2*. A phylogenetic analysis shows that the Burmese python gene LOC103050242 is more closely related to the brown anole LOC132761499 and coelacanth *gch2* than to Burmese python *gch1* (Figure S2). Synteny analysis further supports the identification of LOC103050242 as *gch2*. In humans, coelacanth, and python, *gch1* is flanked by *samd4a*, while in coelacanth and python, *gch2* (LOC103050242 in python) is flanked by *pigf* and *cript* (Figure S3).

## DISCUSSION

Captive-bred ball pythons have emerged as a valuable model system for studying pigmentation biology, offering unique opportunities to explore the genetics of colouration in reptiles (Brown et al. 2022; Dao et al. 2023; Garcia-Elfring et al. 2023; Lederer et al. 2023). In this study, we focused on an axanthic line of ball pythons to investigate the genetic basis of yellow pigmentation in these snakes. While yellow and red hues in vertebrate integument can arise from dietary carotenoids or endogenous pigments, the carnivorous diet of snakes, combined with the yellow colouration observed in wildtype ball pythons, led us to hypothesize that endogenous pigments are the primary source of yellow colouration in this species. Given the normal melanin pattern seen in axanthic ball pythons, we speculated that any candidate mutation would likely be found in a gene associated with pigment biosynthesis rather than in one related to xanthophore cell-fate specification.

Our analysis identified a mutation in *gch2*, a gene known as *ribA* in bacteria, including *E. coli*, where it was first isolated and extensively studied (Richter et al. 1993; Averianova et al. 2020). *Gch2* plays a critical role in the biosynthesis of riboflavin (vitamin B2) and the production of flavin nucleotide coenzymes, which are vital for numerous enzymatic reactions (Bacher et al. 2000; Ren et al. 2005). Unlike animals, bacteria can synthesize riboflavin de novo through an ancestrally conserved pathway, like that found in plants and fungi (Anam et al. 2020; Averianova et al. 2020; Nasuno et al. 2022; Tian et al. 2022). Bacterial studies have shown that *gch2* catalyzes the conversion of GTP into 2,5-diamino-6-ribosyl-amino-4(3H)-pyrimidinedione 5′-phosphate (DARPP), releasing formate and pyrophosphate as byproducts (Averianova et al. 2020; Klein et al. 2023). In animals, although *gch2* can catalyze the GTP to DARPP reaction, its physiological role is considered non-essential since the riboflavin pathway is not fully functional.

The precise function of *gch2* in animals remains less clear compared to that of *gch1* (*GTP cyclohydrolase I*), which is essential for the synthesis of tetrahydrobiopterin (BH4), a cofactor for enzymes involved in neurotransmitter production (e.g., dopamine, serotonin) and nitric oxide synthesis. It also serves as a marker for the xanthophore lineage (Thöny et al. 2000; Ziegler et al. 2000; Pelletier et al. 2001; Nagao et al. 2014; Hamied et al. 2020; Wu et al. 2022). In humans, mutations in *gch1* lead to severe metabolic and neurological disorders (Nagatsu Ichinose, 1996; Longo, 2009; Fanet et al. 2021; Himmelreich et al. 2021), such as phenylketonuria (Trujillano et al. 2017) and DOPA-responsive dystonia (Hirano and Ueno 1999; Yoshino et al. 2018; Kostić et al. 2020; Bradley et al. 2021). *Gch1* and BH4 are also crucial for embryonic development, with homozygous mutations being lethal in mice and zebrafish (Douglas et al. 2015; Larbalestier et al. 2022).

Although *gch2* and *gch1* share functional similarities, such as guanine ring opening, formate release, and zinc binding, the reactions they catalyze differ in significant ways (Auerbach et al. 2000; Kaiser et al. 2002; Ren et al. 2005). The role of *gch2* in vertebrate pigmentation is less well understood, but it has been identified as an early marker for xanthophores (Parichy et al. 2000; Miyadai et al. 2023). In zebrafish, *gch2* is essential for larval xanthophore pigmentation but not in adults (Lister, 2019). Conversely, research in flounder suggests that *gch2* is necessary for adult melanophore and xanthophore differentiation (Watanabe et al. 2008).

We discovered a nonsense mutation at residue 174 (p.Arg174*), likely resulting in a non-functional protein, occurring 16 residues upstream of the conserved histidine believed to be critical for producing DARPP from GTP (Yadav et al. 2020). Our findings indicate that xanthophore pigmentation in ball pythons arises from the riboflavin biosynthesis pathway, rather than the biopterin pathway. Additionally, since loss-of-function mutations in *gch1* often lead to severe developmental defects, the involvement of *gch2*—rather than *gch1*—is consistent with the absence of pleiotropic effects in axanthic ball pythons.

The extensive phenotypic variation in ball python colour morphs and the wide range of colour combinations breeders have produced in captivity provide an opportunity to explore and test hypotheses regarding the mechanisms behind pattern formation. Recent studies suggest that all ball pythons lack iridophores, one of the three chromatophore types (Tzika 2024). This makes ball pythons natural analogs to Shd mutant zebrafish, which also lack iridophores (Frohnhöfer et al., 2013). In addition to axanthism, ball python colour morphs can exhibit xanthism, which is characterized by an increase in yellow brightness (possibly due to melanin changes) or in the distribution of yellow pigment (e.g., the ‘Enchi’ morph). Many ball python colour morphs that seem to affect melanophore migration (e.g., Tri-Stripe, which shows altered melanin patterns) display xanthophore pigmentation in regions typically covered by dark melanin. This non-overlap of black and yellow pigments suggests that xanthophores and melanophores repel each other, as observed in zebrafish (Frohnhöfer et al., 2013).

In zebrafish double mutants (Shd;Pfe), where two of the three chromatophore types (e.g., iridophores and xanthophores) are absent, the remaining chromatophore type (e.g., melanophores) uniformly covers the body, uninhibited by cell-cell interactions. Similarly, we hypothesize that the genetic basis of all-black melanic ball python morphs, such as ‘Cinnamon’ (Figure 1H), is likely due to mutations in genes essential for xanthophore development, allowing melanophores to spread across the body. This contrasts with mutations in melanin production genes, like agouti, which are more common in mammals (e.g., Sasamori et al. 2017; Reissmann et al. 2020). Further support for a repulsive interaction between xanthophores and melanophores comes from the Albino form of the Cinnamon morph (Figure 1I). Instead of revealing an underlying pterin pigment pattern, which might be expected if melanism was driven by excessive melanin production, the ‘Cinnamon Albino’ phenotype presents as an entirely white ball python with no pterin pigmentation (Figure 1I).

Other recessive lines of axanthism exist in captive-bred ball pythons, such as the ‘MJ’ line. Notably, VPI Axanthic and other axanthism lines are not allelic with one another, meaning crosses between different lines result in double heterozygotes with wildtype coloration, suggesting distinct genetic causes for each axanthism lineage. These various axanthism lines, when combined with other color variants (Figure 1D), provide valuable models to further explore the pterin pigment biosynthesis pathway and identify new axanthism-causing genes in reptiles. Consistent with our findings, we demonstrated in the VPI lineage that pterin pigment metabolism, rather than carotenoid metabolism, is associated with yellow coloration in ball pythons.

In summary, our study highlights a significant link between *gch2* and axanthism in ball pythons, suggesting that the initial stages of the ancestral riboflavin pathway are crucial for yellow pigmentation. Further research is necessary to identify the specific xanthophore pigments involved in ball python pigmentation. Nonetheless, our findings enhance the understanding of the genetics of pigmentation in reptiles.

## CONCLUSION

We used ball python colour morphs to study the genetic basis of xanthophore pigmentation in snakes. We found axanthism is associated with a key gene in the biopterin biosynthesis pathway, providing the first evidence that pterins, not carotenoids, produce the yellow pigment observed in ball pythons. To our knowledge, *gch1* is the first axanthism-associated gene identified in vertebrates. Our results provide new research avenues for the study of axanthism and xanthophore pigmentation in snakes and highlights the utility of using captive-bred ball pythons in pigmentation research.

## Acknowledgments

We thank Douglas Menke and Benjamin Wortman for discussions on gene phylogeny and synteny. This work was supported by an NSERC Discovery Grant and Canada Research Chair to RDHB and APH. We thank Mutation Creation, T Dot Exotics, The Ball Room Canada, Ball Python Genetics Canada, and Designing Morphs for supplying samples.

## Data accessibility

The data that support the findings of this manuscript have been uploaded to the NCBI Sequence Read Archive (SRA) with the accession PRJNA1122856.

## Contributions

AGE, APH, and RDHB conceived and designed the study. HR collected samples. AGE and JMA performed the molecular work and AGE the bioinformatics and data analyses. HR performed PCR and Sanger sequencing analysis. AGE wrote the manuscript with input from APH, HR, JMA, and RDHB.

