## Supplementary Figures for "*GTP cyclohydrolase II* (*gch2*) and axanthism in ball pythons: a new vertebrate model for pterin-based pigmentation"

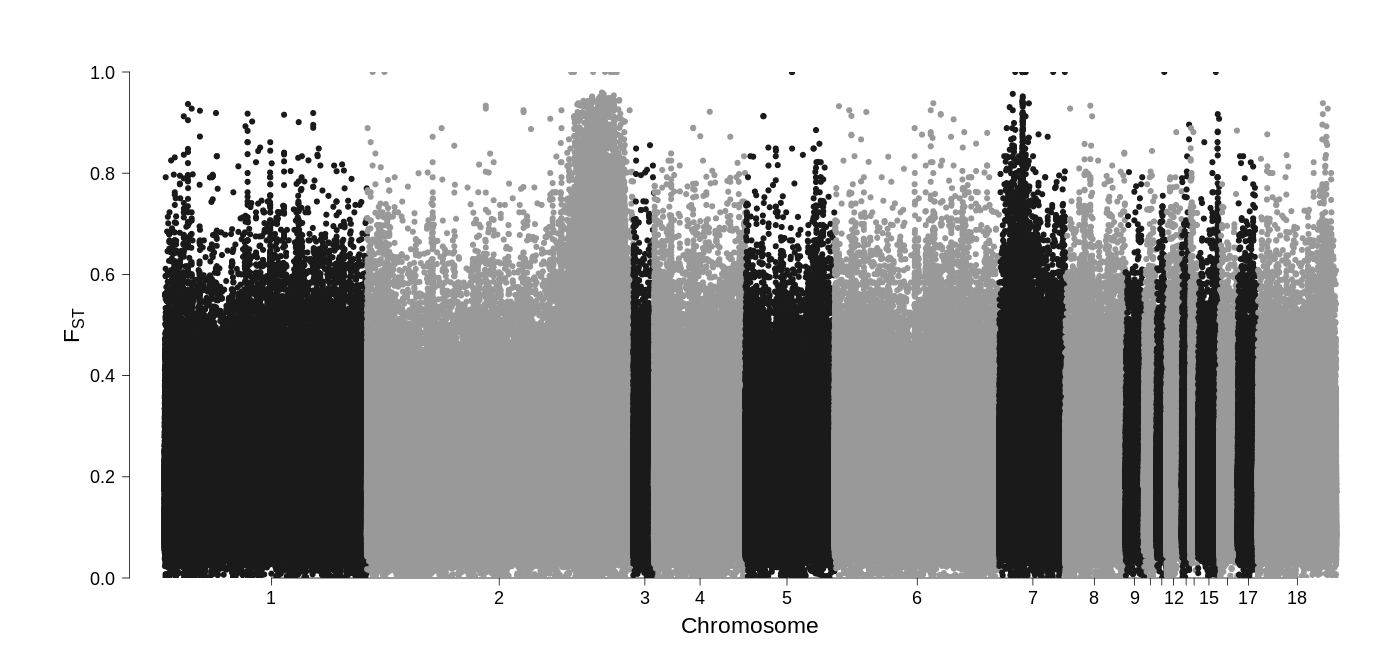

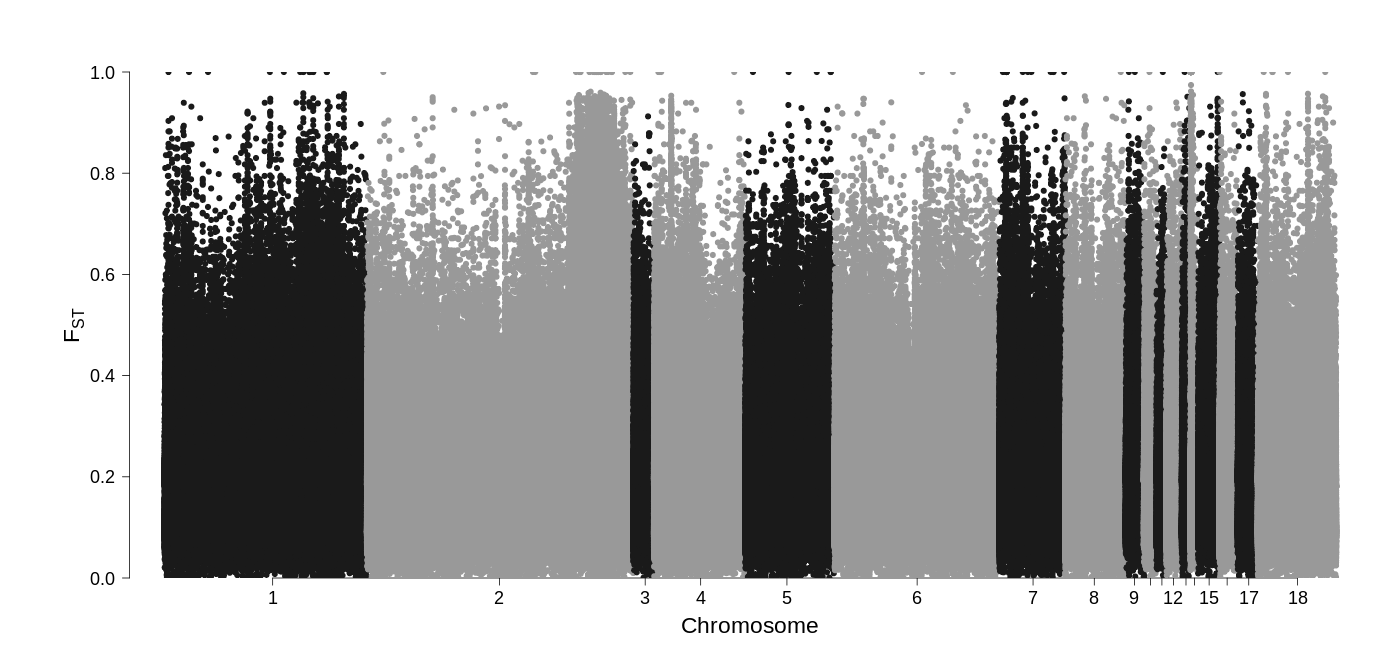

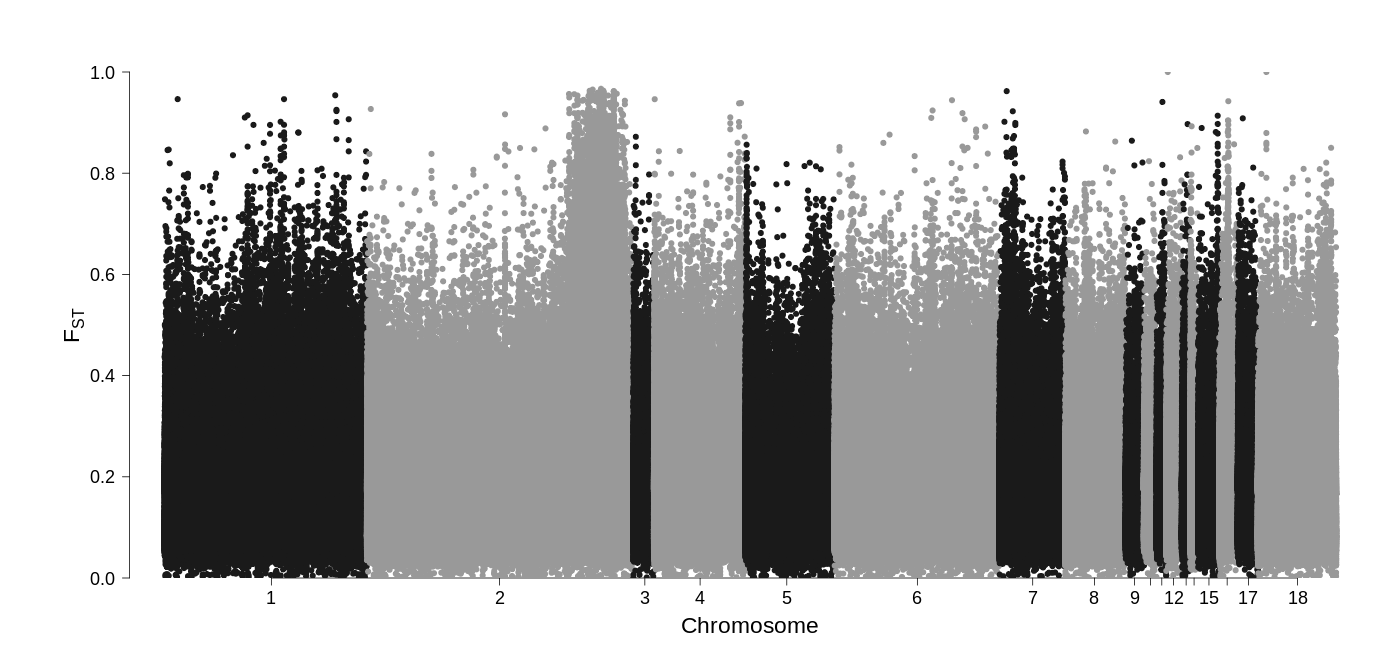

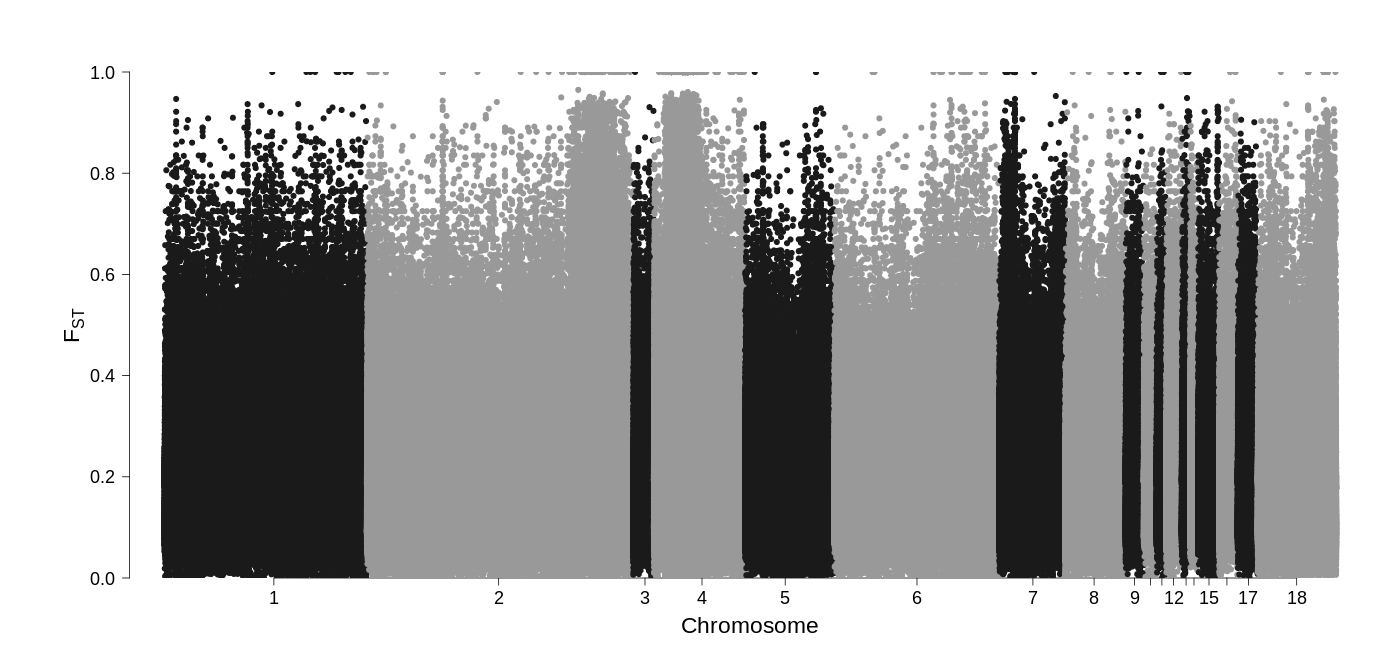

Figure S1. Manhattan plots of genome-wide differentiation (F_ST_) between the axanthic (i.e., case) and control pools (from the top: ‘Spider’, n=26; ‘Enchi’, n=7; ‘Ivory’, n=15; ‘Lavender’, n=14). The region on chromosome 2 consistently differentiated in case and control comparisons contains the candidate gene LOC103050242 (*gch2*).

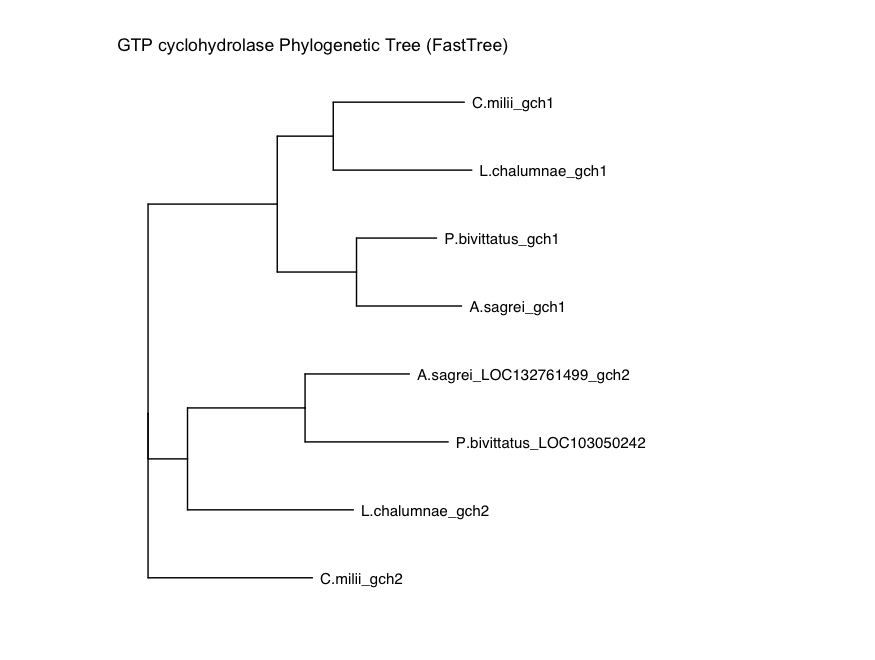

Figure S2. Phylogenetic tree of *gch1*, *gch2*, and LOC103050242 across different vertebrates: Burmese python (*Python bivittatus*), brown anole (*Anolis sagrei*), coelacanth (*Latimeria chalumnae*), and elephant shark (*Callorhinchus milii*).

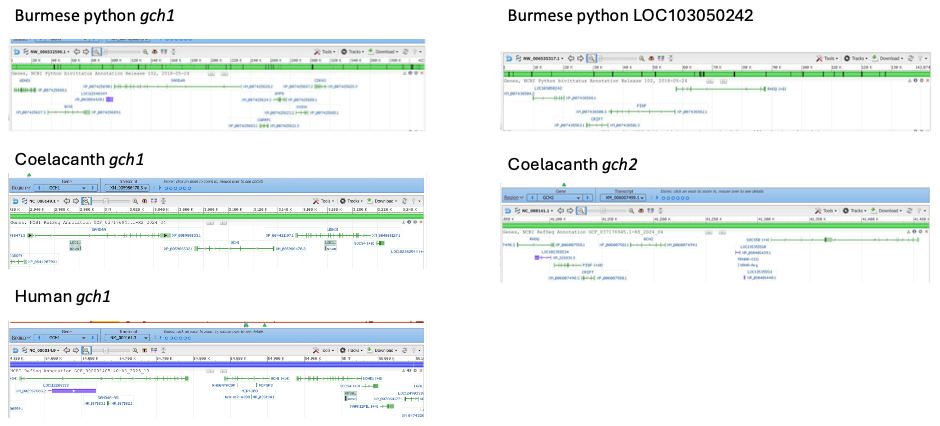

Figure S3. Synteny analysis for *gch1*, *gch2*, and LOC103050242. In humans, coelacanth, and python, *gch1* is flanked by *samd4a*. Coelacanth *gch2* and Python LOC103050242 are flanked by *pigf* and *cript*.
